# Not all smokers are alike: The hidden cost of sustained attention during nicotine abstinence

**DOI:** 10.1101/2021.07.22.453142

**Authors:** Harshawardhan U. Deshpande, John R. Fedota, Juan Castillo, Betty Jo Salmeron, Thomas J. Ross, Elliot A. Stein

**Affiliations:** Neuroimaging Research Branch, National Institute on Drug Abuse-Intramural Research Program, National Institutes of Health, Baltimore, MD, USA; Department of Psychology, Harvard University, Cambridge, MA, USA; Behavioral and Cognitive Neuroscience Branch, Division of Neuroscience Behavior, National Institute on Drug Abuse-Intramural Research Program, National Institutes of Health, Bethesda, MD, USA

**Author notes:** **Correspondence to:** Elliot A. Stein, Neuroimaging Research Branch, National Institute on Drug Abuse-Intramural Research Program, National Institutes of Health, Baltimore, MD, USA, John R. Fedota, Behavioral and Cognitive Neuroscience Branch, Division of Neuroscience Behavior, National Institute on Drug Abuse-Intramural Research Program, National Institutes of Health, Bethesda, MD, USA. These authors have contributed equally to the work (Harshawardhan U. Deshpande, John R. Fedota).

**Keywords:** nicotine use disorder, fMRI, functional connectivity, attentional control

## Abstract

**Background:** Nicotine Withdrawal Syndrome (NWS)-associated cognitive deficits are heterogeneous, suggesting underlying endophenotypic subgroups. We identified smoker subgroups based on response accuracy during a cognitively demanding Parametric Flanker Task (PFT) and characterized their distinct neuroimaging endophenotypes using a nicotine state manipulation (sated, abstinent).

**Methods:** Forty-five smokers completed the 25-min PFT in two fMRI sessions (nicotine sated, abstinent). Task-evoked NWS-associated errors of omission (EOm), brain activity, underlying functional connectivity (FC), and brain-behavior correlations between subgroups were assessed.

**Results:** Based on their response accuracy in the high demand PFT condition, smokers split into high (HTP, n=21) and low task performer (LTP, n=24) subgroups. Behaviorally, HTPs showed greater response accuracy independent of nicotine state and greater vulnerability to abstinence-induced EOm. HTPs showed greater BOLD responses in attentional control brain regions for the [correct responses (–) errors of commission] PFT contrast across states. A whole-brain FC analysis with these subgroup-derived regions as seeds revealed two circuits: L Precentral : R Insula and L Insula : R Occipital, with abstinence-induced FC strength increases only in HTPs. Finally, abstinence-induced brain (FC) and behavior (EOm) differences were positively correlated for HTPs in a L Precentral : R Orbitofrontal cortical circuit.

**Conclusion:** We used a cognitive stressor (PFT) to fractionate smokers into two subgroups (HTP/LTP). Only the HTPs demonstrated sustained attention deficits during nicotine abstinence, a stressor in dependent smokers. Unpacking underlying smoker heterogeneity with this ‘dual stressor’ approach revealed distinct smoker subgroups with differential attention deficit responses to withdrawal that could be novel targets for therapeutic interventions to improve cessation outcomes.

## 1. Introduction

Following acute smoking abstinence, most smokers exhibit components of the Nicotine Withdrawal Syndrome (NWS), manifest as a set of aversive affective, somatic, and cognitive disruptions peaking in the initial days of a quit attempt (1). The timing and severity of the NWS symptoms are salient to and predictive of long-term smoking cessation, with more symptoms generally associated with lower success (2,3). Indeed, the NWS often dissuades smokers from continued abstinence through negative reinforcement, thus promoting relapse to alleviate the withdrawal state (4–6).

The clinical symptoms of the NWS vary greatly in their duration, intensity, and phenomenology across individuals (7–9). Indeed, only some smokers even report abstinent-induced overt cravings and anxiety (10). Moreover, the efficacy of current FDA approved cessation aids such as bupropion (11), varenicline and nicotine replacement therapy (NRT(12)) is quite variable, suggesting heterogeneity in the smoking phenotype. Further, genetically informed biomarkers such as the nicotine metabolite ratio induced by variations in the hepatic enzyme cytochrome P450 (CYP2A6(13)) predict success with NRT and lend further credence to the underlying population diversity. Taken together, these studies suggest the existence of smoker subgroups that relate to the diversity of NWS symptom duration and intensity and variable cessation treatment outcomes (14), suggesting that the current high treatment failure rate may be improved by tailoring treatments to *a priori* identified smoker subgroups.

One potential way to differentiate subgroups of smokers is via characterization of NWS-induced cognitive disruptions. Chronic stress from abstinence-induced cognitive disruptions likely results in a maladaptive response known as allostatic load (15–17). For instance, longstanding elevated activation of the brain’s stress systems reduces a smoker’s ability to respond when presented with additional extrinsic stressors, thus enhancing the reinforcing effects of nicotine during abstinence, increasing relapse vulnerability (5,18).

Moreover, cognitively demanding tasks can produce autonomic stress-like responses (19,20). During nicotine withdrawal, smokers often report impairments in cognitive control, including decrements in performance on tasks reflecting sustained attention (21–23), response inhibition (24), and working memory (25–27), with these cognitive deficits predictive of smoking relapse (28,29). Previous studies have observed a parametric response to task demand in abstinent smokers, especially at high levels of task difficulty (25,30).

Acute nicotine abstinence is also a stressful period for smokers (31,32). For example, Sutherland and colleagues (33) found that in abstinent (vs. sated) smokers, the resting state functional connectivity (rsFC) strength in an amygdala-insula-Default-Mode Network (DMN) circuit was downregulated following NRT administration, suggesting relief of the reported subjective withdrawal state. Fedota et al. (34) found that dorsal and posterior insular rsFC circuits with the DMN and the Salience Network (SN) are enhanced during abstinence (vs. satiety), while a ventral insular rsFC connection to the Executive Control Network (ECN) was reduced, and time-varying FC changes show reduced temporal flexibility and lower network spatiotemporal diversity between abstinence and satiety (35).

Together, these observations demonstrate the utility of cognitive demand and abstinence as stressors that reveal changes in rsFC in smokers. Experimentally challenging neural regulatory mechanisms in smokers via an imposed ‘stress test’ in the form of an allostatic challenge should thus expose otherwise unobserved maladaptive processes and identify smoker subgroups that will differentially respond to an extrinsic stressor. In the current study, we leveraged a ‘two-level stress test’ of a cognitively demanding parametric flanker task (PFT) and acute abstinence as simultaneous allostatic stressors to enhance the variability in maladaptive processes and neurobiological mechanisms to: a) fractionate smokers into distinct endophenotypic subgroups and b) characterize the differential behavioral and neurobiological responses of these subgroups to nicotine abstinence. Using a within-subjects design, we hypothesized that subgroups would be identified based on differential cognitive performance and that the subgroups would show further differences in task-evoked NWS-related cognitive behaviors (lapses of attention), task-evoked brain activation, and underlying task-related FC patterns.

## 2. Methods

### 2.1. Participants

Fifty-nine right-handed smokers were recruited into the study. Eleven were excluded during final analyses (two for incomplete behavioral data, two for scanning errors, two for abnormal MRI and five for excessive head motion). Overall demographics for the remaining forty-five participants are described in Table 1. Previously published studies include a subset of these participants (34,36).

**Table 1.**
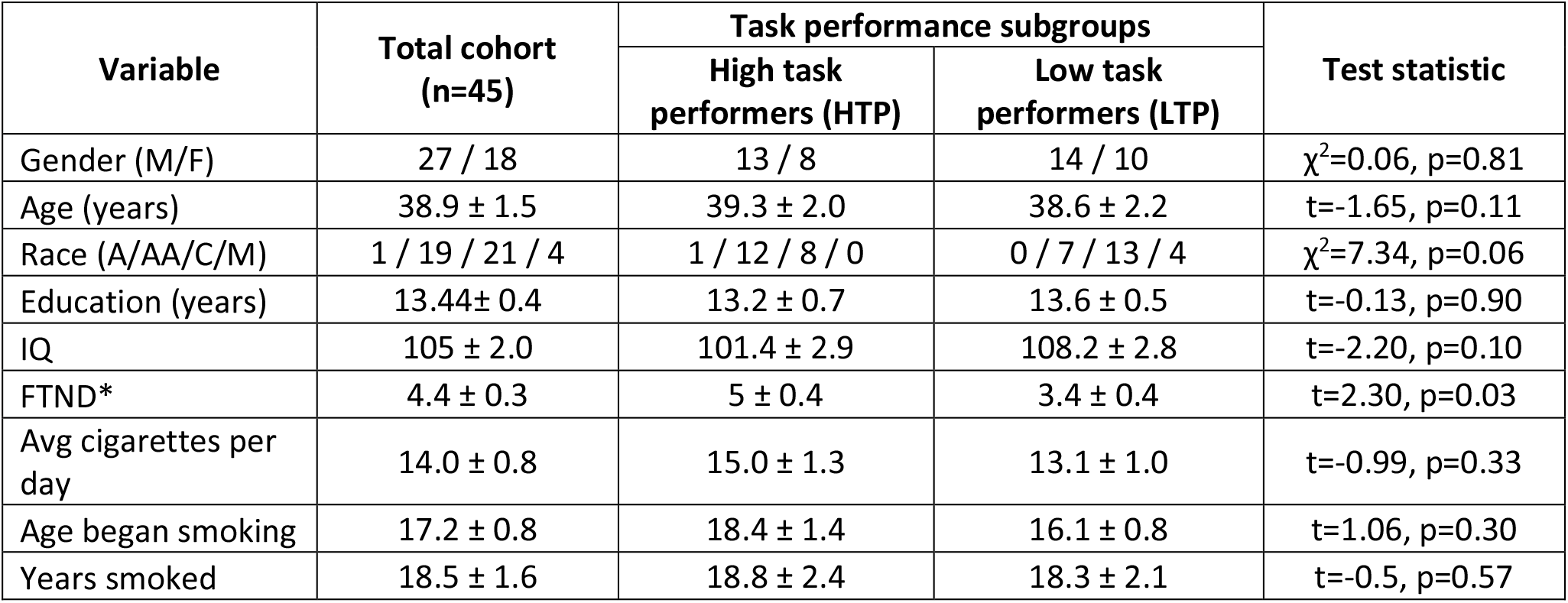
Demographics for the whole cohort (n=45) of smokers and both SUBGROUPs (n=21 for HTP and n=24 for LTP based on adjusted accuracy on the high DEMAND condition of the Parametric Flanker Task). A, Asian; AA, African American; C, Caucasian; M, Mixed; IQ, Intelligence Quotient; FTND, Fagerström Test of Nicotine Dependence; * denotes a significant group difference. Values represent mean ± standard error.

### 2.2. Experimental Design

In a longitudinal within-subjects design, participants completed two MR scanning sessions – one during sated smoking and another after ∼48 hours of biochemically verified nicotine abstinence. The order of the two scan sessions was fixed as these data are part of a larger ongoing smoking cessation protocol (clinicaltrials.gov identifier: NCT01867411). The sated scan preceded the abstinence scan by an average of 75 days (median 28 days). See Fig. 1 for experimental design overview.

**Figure 1.**
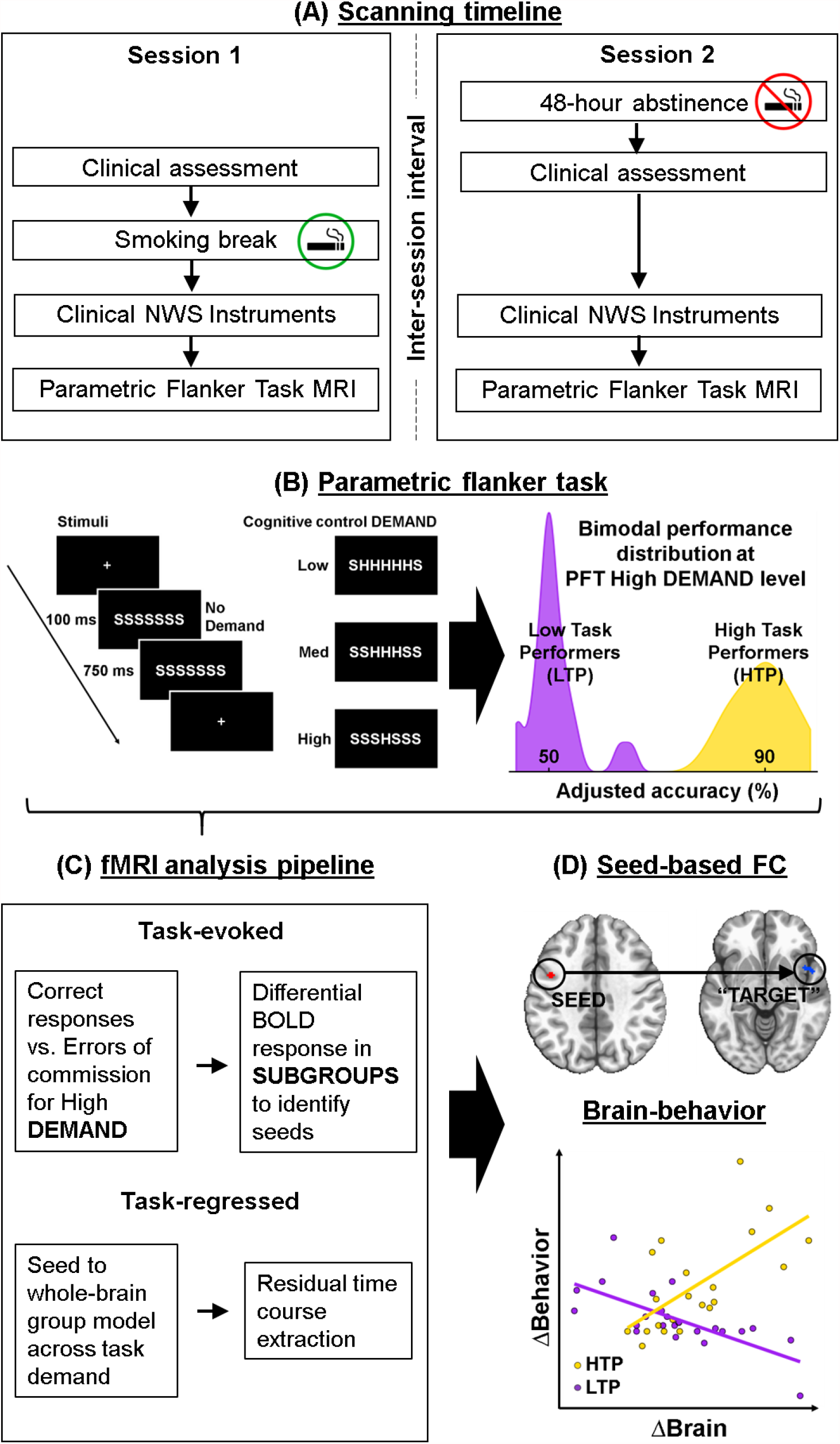
Study experimental design, data analysis pipelines and key findings. **(A)** In a within-subjects design with two scanning sessions, participants were nicotine sated (session 1) or ∼48 hours abstinent (session 2). The inter-session interval averaged at 75 days (median 28 days). **(B)** During both the MRI scans, participants performed a 25-minute Parametric Flanker Task (PFT). Cognitive control DEMAND is modulated via the number of conflicting stimuli flanking the target stimuli (no, low, medium, or high DEMAND). Based on the adjusted accuracy at the high DEMAND, participants were divided into two SUBGROUPs: Low task performers (LTP) and High task performers (HTP). **(C)** The task-evoked pipeline was used to identify brain regions showing greater differential response in high DEMAND for the [correct responses (–) errors of commission] for HTPs vs. LTPs. Using the task-regressed functional connectivity (FC) pipeline, these regions were used in a seed-based FC analysis **(D)** to identify seed-“target” dyads with FC SUBGROUP differences between STATEs. Finally, SUBGROUP STATE differences in FC and Errors of Omission (EOm) were related.

### 2.2.1. Behavioral and subjective measures

Participants were scanned while completing a PFT. Data were collected on two Siemens scanners (Trio and Prisma). The PFT was modified from the classic Eriksen flanker task (36,37) to instantiate varying levels of DEMAND for cognitive control on a trial-by-trial basis (Fig. 1B). All stimuli were presented using E-Prime software (Psychology Software Tools, Sharpsburg, PA).

Subjective ratings of withdrawal (Wisconsin Smoking Withdrawal Scale, WSWS (38)), craving (Tobacco Craving Questionnaire Short Form, TCQ-SF (39)) and affect (Positive and Negative Affect Schedule, PANAS (39,40)) were assessed prior to each scanning session.

See the Supplement for details of the PFT, subjective measurements and MRI acquisition parameters.

### 2.3. Data Analysis

#### 2.3.1. Behavioral Task and subjective measures

Behavioral effects of smoking: During PFT performance, STATE (satiety/abstinence), and demand for cognitive control: DEMAND (none/low/medium/high) were quantified via correct response speed (*Speed*= 1/Reaction Time), coefficient of variation of correct response speeds (*SpdCV*=std(SpeedCorrectTrials)/mean(SpeedCorrectTrials)), trial adjusted accuracy (*Accuracy*=Correct Trials)/(Correct Trials + Errors of Commission(ECo)), and Errors of Omission (EOm) – a measure of lapses in attention. Performance accuracy in the high DEMAND condition identified SUBGROUPs of participants (High Task Performers/Low Task Performers, HTP/LTP). All subsequent behavioral and neuroimaging analyses accounted for these SUBGROUPs. All analyses were conducted using R.

*Speed, SpdCV*, and *Accuracy* were submitted to mixed-design analysis of variance (ANOVA) with STATE and DEMAND as within-subjects factors, and SUBGROUP as a between-subjects factor. Significant interactions were evaluated via post-hoc paired and unpaired t-tests where appropriate. ΔEOm were calculated as a function of STATE (abstinence [-] sated) prior to examination of between-subject SUBGROUP differences via Kruskal-Wallis one-way ANOVA. Significant non-parametric effects were evaluated via post-hoc Wilcoxon rank-sum test.

Assumptions of normality in the subjective measures (WSWS, TCQ, PANAS) were evaluated following calculation of the difference as a function of STATE (Δ Score; abstinence [-] sated). STATE effects were calculated via one-sample t-tests comparing Δ Score against null hypotheses of 0; between-subject differences in SUBGROUP identity were evaluated via independent t-tests. Where appropriate, equivalent nonparametric statistical tests were utilized.

#### 2.3.2. Imaging

The PFT fMRI data were processed and analyzed using AFNI(41) in two parallel pipelines: task-evoked activation and task-regressed functional connectivity. The task-evoked pipeline modeled differences between SUBGROUPS (HTP/LTP) in task-evoked responses during the high DEMAND condition of the PFT. In contrast, since lapses of attention (EOm) were observed across DEMAND conditions, the PFT-evoked responses were regressed out in a task-regressed pipeline.

##### Task-evoked Pipeline

###### Individual level analysis

After preprocessing (see Supplement), an event-related analysis of the PFT data was performed using a voxel-wise multiple regression analysis with regressors expressed as a delta function convolved with a standard hemodynamic response function (SPM gamma variate basis function) and its temporal derivative (*AFNI: 3dDeconvolve*). Regressors included DEMAND (none, low, medium, high) for both correct and error responses (eight total regressors) and six head motion parameters. A voxel-wise average amplitude change equal to the percentage change from baseline (β) was calculated per participant and regressor. The design matrix obtained was applied to the concatenated normalized time series (*AFNI: 3dREMLfit*) to obtain the beta + statistics dataset with restricted maximum likelihood estimation and the dataset for the REML residuals. The minimum voxel cluster size for all whole-brain analyses was determined (*AFNI: 3dFWHMx, 3dClustSim*) using a two-component measure of the spatial autocorrelation of the preprocessed data (42).

###### Group level analysis

To identify task activation differences to serve as FC seeds, a multivariate model approach (*AFNI: 3dMVM*) was used with a SUBGROUP contrast (HTP vs. LTP) of correct trials vs. error trials at the high DEMAND condition. Factors in the model were STATE (satiety, abstinence) and SUBGROUP (HTP, LTP) with scanner (Trio, Prisma) and ΔFD head motion (abstinent [-] sated) as covariates. A conservative voxelwise threshold of p=0.0001 applied on the SUBGROUP difference for the high DEMAND [correct responses (–) errors of commission] contrast identified 19 spatially specific clusters (minimum k=23 voxels; FWE of α≤0.05). These clusters were then used in a seed-based FC analysis on the task-regressed data to identify SUBGROUP FC differences to relate with EOm.

##### Task-regressed Pipeline

The significant clusters of PFT-evoked activation showing SUBGROUP differences obtained from the above *task-evoked pipeline* were used as regions-of-interest (ROIs) in a seed-based FC analysis to identify FC SUBGROUP differences and their relationship with lapses of attention (EOm) across all DEMAND conditions. As using task-evoked data to perform seed-based FC analyses can produce false positives and inflate the FC estimates (43), seed-based FC analyses were conducted after regressing out the PFT and modeling the hemodynamic response function (HRF) using a finite impulse response model. The HRF was modeled as a set of tent functions (8 parameter TENT function for 14 time points, *AFNI: 3dDeconvolve, 3dREMLfit*). The residual time series was used as a proxy for resting-state data to characterize differences between SUBGROUP FC and the relationship of SUBGROUP FC to EOm. Data were band pass filtered between 0.001 Hz to 0.25 Hz (*AFNI: 3dTproject*).

##### Seed-based FC

While the SUBGROUPs were revealed by behavioral *Accuracy* on the high DEMAND PFT condition, a SUBGROUP*STATE effect on *Errors of Omission* was observed across all task conditions. To characterize the network interactions underlying this decrease in sustained attention in the HTP SUBGROUP, a multivariate model (*AFNI: 3dMVM*) was created for each of the observed 19 seeds; whole-brain FC was examined for a SUBGROUP main effect and a SUBGROUP*STATE interaction. A voxelwise threshold of p=0.001 (k= 69 voxels; FWE of α≤0.05) was used to identify clusters with a SUBGROUP*STATE interaction from each seed to whole brain. Pairs of regions with significant FC interaction after voxelwise and cluster thresholding are denoted as *dyads*, with the task activation difference pole denoted as the “seed” and the differential FC denoted as the “target” pole. The “target” pole was subsequently used as a seed in a second whole-brain FC analysis to identify regions with which it had differential functional connections (e.g. Sutherland et al. 2013a(33)) to define larger functional circuits differing between HTPs and LTPs.

##### Relating STATE differences in functional connectivity (ΔFC) with lapses of attention (ΔEOm)

To relate FC STATE differences with changes in behavior relevant to EOm, a multiple linear regression analysis was modeled (*AFNI: 3dRegAna*). The model included ΔEOm, SUBGROUP membership and a SUBGROUP*ΔEOm interaction term. ΔFD was included as a covariate to help account for residual motion. Since six separate regression analyses were conducted, a threshold of p=0.001 (k=100 voxels; FWE of α≤ (0.05/6) was used to correct for multiple comparisons.

## 3. Results

### 3.1 Cohort behavioral and subjective measures

There was a main effect of DEMAND with lower Speed, SpdCV and Accuracy for the high DEMAND and a main effect of STATE with Accuracy lower and EOm higher during abstinence. As expected, subjective ratings of withdrawal, craving and affect also showed a main effect of STATE. See Supplement for detailed behavioral/subjective results.

### 3.2 Subgroup behavioral and subjective differences

A clear dichotomy was observed in *Accuracy* in the high DEMAND PFT condition across nicotine STATE, which defined two SUBGROUPs: High task performers (HTP, N=21) with 88.68% (± 5.19SD) *Accuracy* and Low task performers (LTP, N=24) with 51.04% (± 4.72SD) *Accuracy*. Table 1 and Fig 2 illustrate clinical instruments, and other behavioral task performance measures for the two SUBGROUPs.

**Figure 2.**
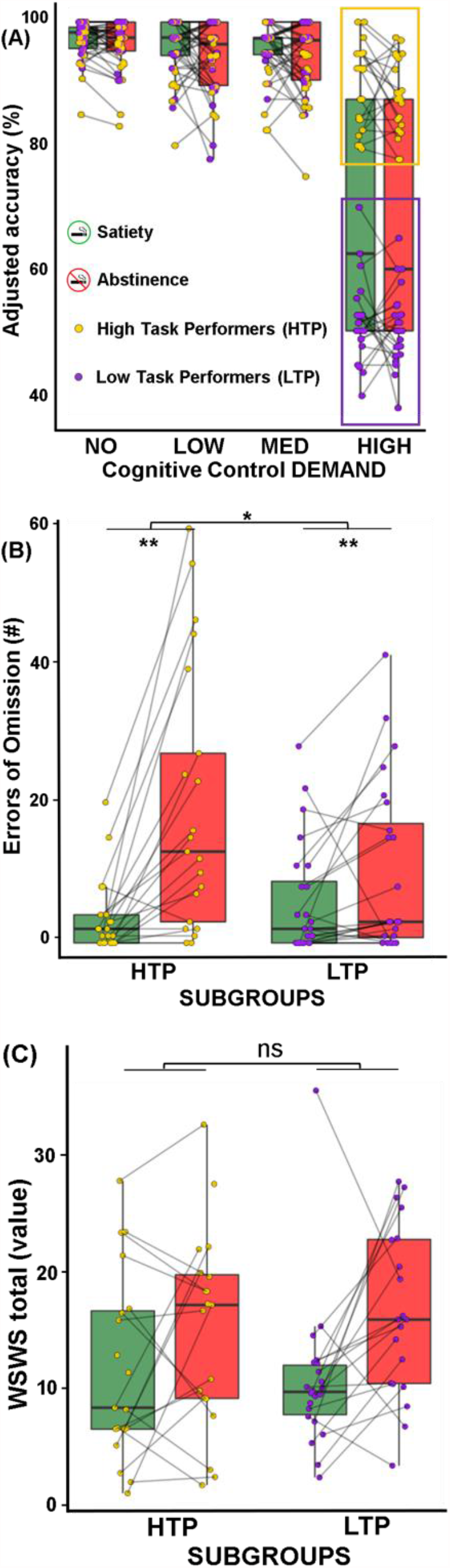
Behavioral performance on the PFT distinguishes cohort into two distinct SUBGROUPs: **(A)** In both satiety (green) and abstinence (red), High task performers (HTP, yellow box, 88.68% ± 5.19 SD) show higher accuracy only in the High DEMAND condition compared to Low task performers (LTP, purple box, 51.04% ± 4.72 SD) for responded trials. **(B)** Across all demand conditions, HTP show a greater increase (Kruskall-Wallis chi-squared = 5.4741, p = 0.01) in the number of Errors of Omission (EOm) during abstinence compared to LTP; HTP: W=378, p=8.76e-06, LTP: W=432, p=9.47e-4. **(C)** There was no significant difference between the HTP and LTP SUBGROUPs for difference in the total Wisconsin Smoking Withdrawal Scale (p=0.29, F=1.17), between satiety and abstinence. Other subjective measures (PANAS, TCQ) also did not show a difference between nicotine STATE for the two SUBGROUPs (see Supplement). HTP: High Task Performers, LTP: Low Task Performers, PANAS: Positive and Negative Affect Scale, TCQ: Tobacco Craving Questionnaire

#### 3.2.1. Demographics

Perhaps counterintuitively, the HTPs had greater nicotine dependence (p=0.03; FTND score for HTPs 5 +/- 0.38SD) compared to the LTPs (3.83 +/- 0.35SD)). No other demographic measures differed significantly between the SUBGROUPs.

#### 3.2.2. Task performance

There was a significant SUBGROUP*STATE ΔEOm effect, with higher attention lapses for the HTPs (Kruskall-Wallis chi-squared = 5.4741, df=1, p = 0.01). Both SUBGROUPs had an increase in EOm in abstinence (HTP: W=378, p=8.76e-06, LTP: W=432, p=9.47e-4). No other SUBGROUP or SUBGROUP*STATE effects were observed.

#### 3.2.3. Subjective measurements

No SUBGROUP differences in subjective measures were found for withdrawal, craving, affect nor any SUBGROUP*STATE interactions. Both groups showed the expected STATE effects (higher negative and lower positive values in clinical instruments).

### 3.3. Neuroimaging

The current analyses focus on neuroimaging SUBGROUP differences. See Supplement Fig. S2 for the overall PFT task map.

#### 3.3.1. Subgroup differences in the high DEMAND condition (Correct – ECo)

Based on SUBGROUP difference in task accuracy, we identified regions of task-evoked differences using the high DEMAND [correct responses (–) errors of commission] contrast trials between SUBGROUPs (Fig. 3A, p-voxel <0.001; p-corrected<0.05) across nicotine STATE. Only the HTP SUBGROUP showed a larger BOLD response for error vs. correct trials in regions including bilateral insula, dorsal ACC, frontoparietal attentional regions and right thalamus (exemplar ROI extract in Fig. 3B). A complete list of activated regions and BOLD signal ROI extracts are in the Supplement (Fig. S3).

**Figure 3.**
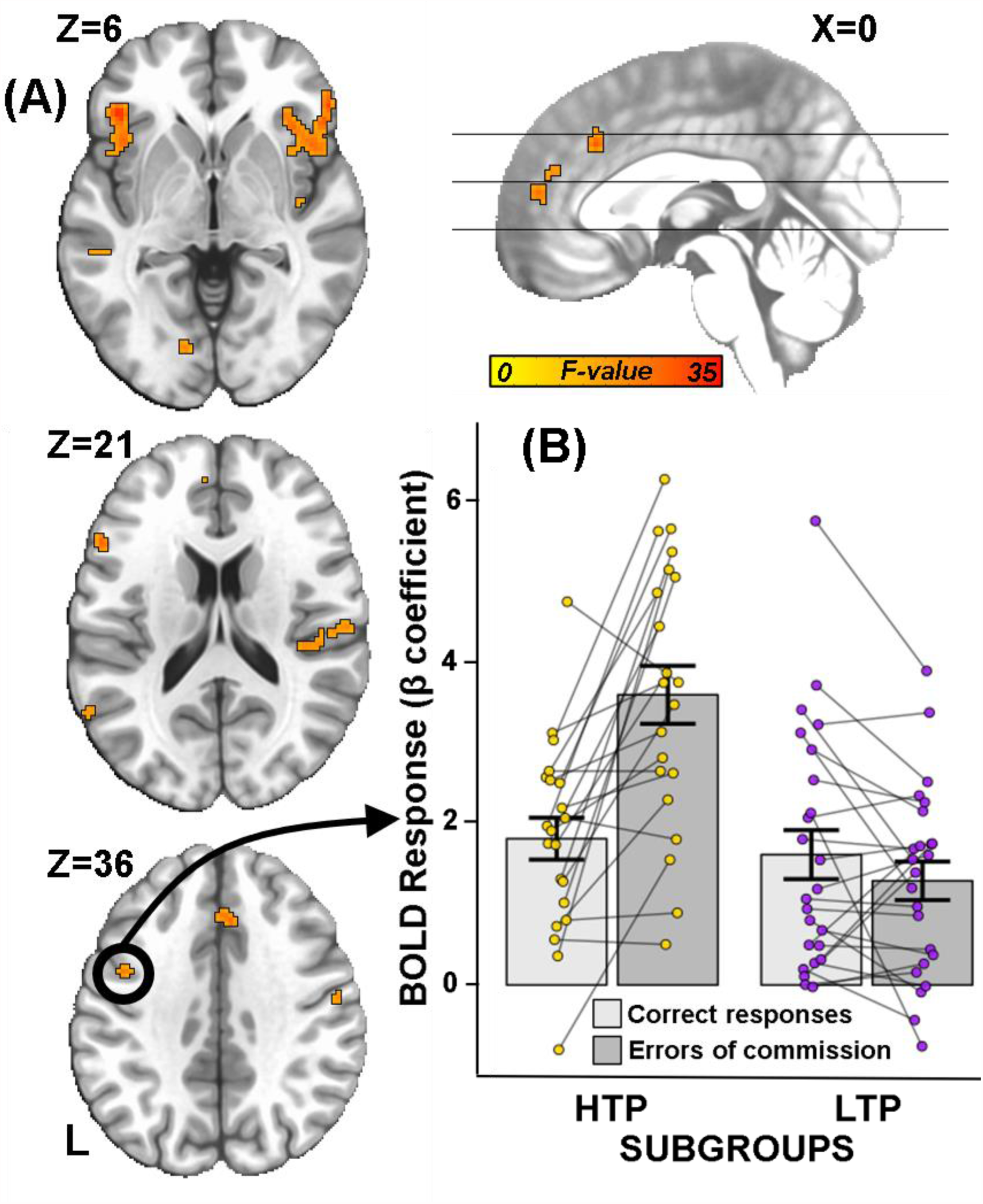
Parametric Flanker Task evoked BOLD response for the high DEMAND responded trials. **(A)** Clusters with differential activation for the HTP compared to the LTP SUBGROUP for the [correct responses (–) errors of commission] trials contrast at the high DEMAND condition across SESSION. These clusters were subsequently used as seeds in a whole-brain seed-based functional connectivity analysis. **(B)**Extracted β coefficients from the encircled cluster (averaged across SESSION) showing greater sensitivity to errors of commission vs. correct responses in the HTP SUBGROUP and no difference between correct responses and errors of commission for the LTP SUBGROUP. For ROI extracts from all clusters see Fig. S3. HTP: High Task Performers, LTP: Low Task Performers, ROI: Region of Interest

#### 3.3.2. Subgroup differences in seed-based FC (task-regressed analyses)

To describe network communication during task performance, the 19 clusters derived from the PFT-evoked differences described above were used as seeds in a whole brain FC analysis. This was done to interrogate the relationship between FC and the behavioral SUBGROUP*STATE effects observed in EOm across all PFT DEMAND conditions. Two of the ROIs employed as seeds showed a SUBGROUP*STATE interaction (Fig. 4A, p-voxel<0.001; p-corrected<0.05). Specifically, the L Precentral (LPre) seed showed a SUBGROUP*STATE interaction with R ventral insula (RvI), while the L posterior insula (LpI) seed showed a SUBGROUP*STATE interaction (Fig. 4A, p-voxel<0.001; p-corrected<0.05) with the R Mid Occipital (RmO) region. These identified circuits (LPre : RvI and LpI : RmO) are denoted as *dyads* 1 and 2, respectively, with one pole of the *dyad* being the seed and the other pole being the functionally connected ‘target’. The LPre : RvI and the LpI : RmO circuits both showed an increased FC (extracted averaged Z-scored FC in Fig. 4B) in the HTPs during abstinence but no significant change for the LTPs, established by separate analyses for each SUBGROUP.

**Figure 4.**
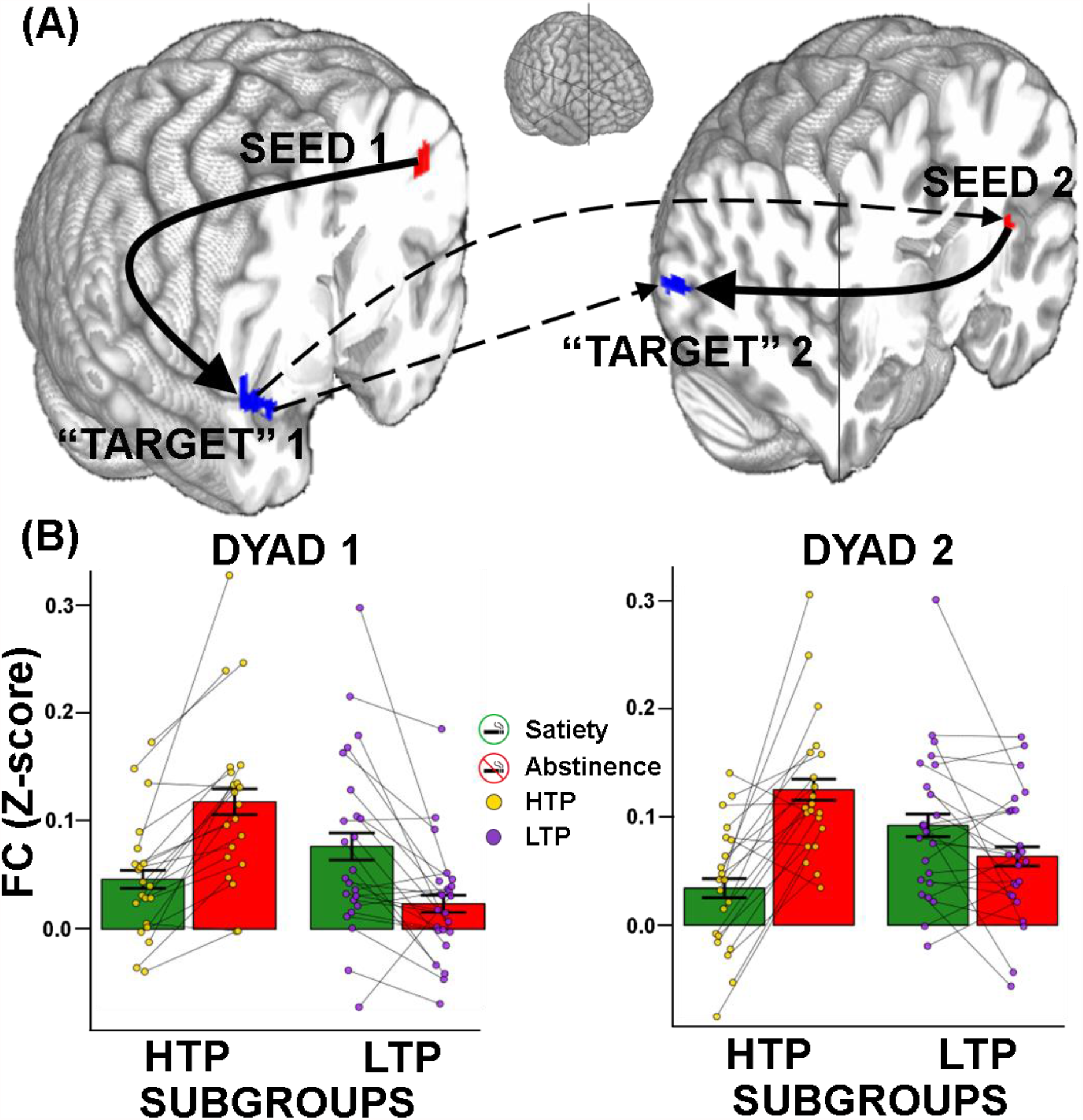
Functional connectivity is higher for HTP during PFT-regressed fMRI: **(A)** *dyad1* with the L Precentral seed (L Pre, red) and the R vent Insula “target” (RvI, blue) and *dyad2* with the L pos Insula seed (LpI, red) and the R mid Occipital “target” (RmO, blue). A “target” in our analysis was defined as a significant cluster arising from a whole-brain seed-based FC analysis. The solid arrows indicate within-*dyad* FC while the dotted arrows indicate between-*dyad* FC. The arrows for seed-“target” and *dyad*-*dyad* FC show do not imply directionality or causality. **(B)** Z-scored FC averages showing increased FC during nicotine abstinence (red bars, SUBGROUP*STATE interaction) within *dyad1* (left, LPre : RvI) and *dyad2* (right, LpI : RmO). Error bars show standard error of the mean. Please see Supplemental table S2 for “seed” and “target” MNI coordinates. L Pre: Left Precentral gyrus, RvI: Right ventral Insula, LpI: Left posterior Insula, RmO: Right middle Occipital

#### 3.3.3. FC coactivation between dyad1 and dyad2

Each of the ‘target’ poles from the initial task FC analyses (dyads 1 and 2) was then used as a seed in a second whole-brain FC analysis to further describe network interaction differences between the SUBGROUPs.

##### *Dyad1* (LPre : RvI)

In a SUBGROUP*STATE interaction, increased FC is observed between the RvI and LpI for the HTPs during abstinence, indicating functional coactivation between the RvI (*dyad1* pole) and the LpI (*dyad2* pole).

##### *Dyad2* (LpI : RmO)

In a SUBGROUP*STATE interaction, increased FC is observed between the RmO and RvI for the HTPs during abstinence, indicating functional coactivation between the RmO (*dyad2* pole) and the RvI (*dyad1* pole).

Thus, the pair of *dyads* are functionally coactivated (Fig. 4A, dotted lines).

#### 3.3.4. Brain-behavior interactions: SUBGROUP FC differences with lapses of attention (EOm)

The relationship between the SUBGROUP FC difference and the key behavioral SUBGROUP*STATE difference, i.e., EOm, was characterized using a multiple linear regression analysis. The seeds of this analysis were derived from the poles of the dyads in the above task-regressed analysis. To test the SUBGROUP*STATE relationship, ΔEOm (abstinent [–] sated behavioral metric) and ΔFC (abstinent [–] sated brain metric) were correlated for each SUBGROUP. Using the LPre in dyad1 as a seed, there was a significant positive relationship between ΔEOm and ΔFC for the HTPs and a negative relationship for the LTPs in the R orbitofrontal cortex (Fig. 5A). Additionally, using the RvI in dyad1 as a seed, there was a positive relationship between ΔEOm and ΔFC for both the HTPs and the LTPs in the L Medial Frontal cortex as a main effect of ΔEOm (Fig. 5B). Similar positive relationships between ΔEOm and ΔFC for both the HTPs and the LTPs were observed using the RmO in dyad2 as a seed in regions including R parahippocampal, L Middle Temporal and R Lingual gyri (main effect of ΔEOm, Supplement Fig. S4).

**Figure 5.**
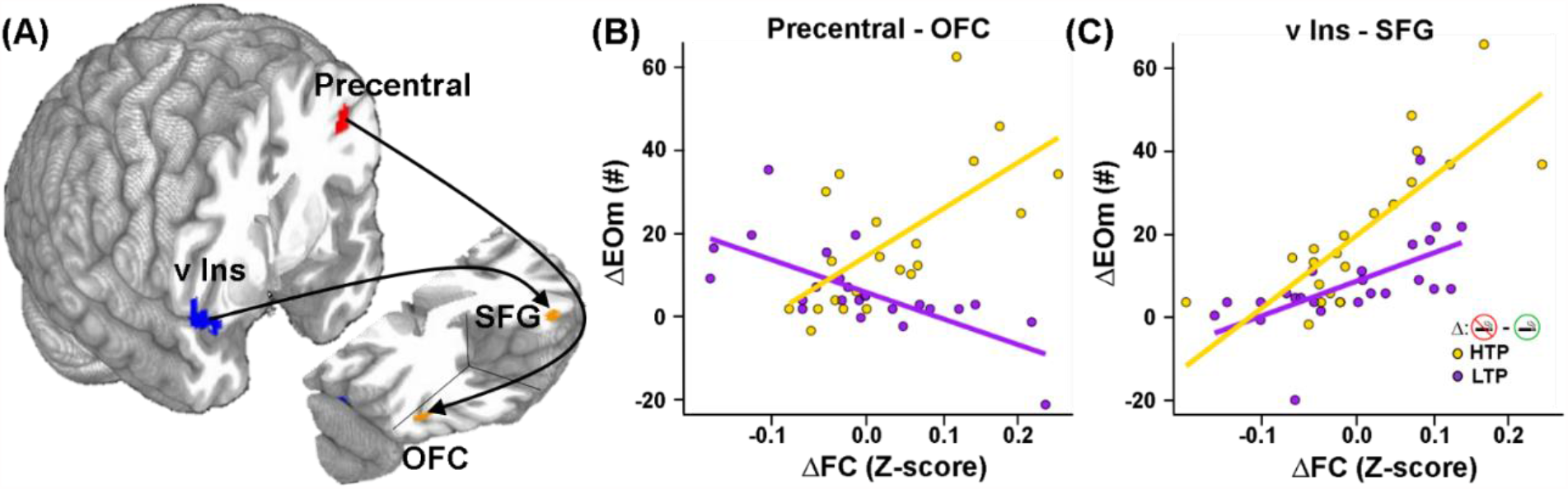
Brain (ΔFC) and behavior (ΔEOm) regression between SUBGROUP STATE differences (abstinence – satiety) using individual Z-scored FC extracts (averaged over “target” ROI) and EOm overall DEMAND levels of the PFT. **(A)**Whole-brain regression analyses starting with seeds at the L Precentral (L Pre, red) and R ventral Insula (RvI, blue) lead to “targets” in the R orbitofrontal cortex (R OFC) and L superior frontal gyrus (L SFG) respectively (orange). The arrows do not imply directionality or causality. **(B)** The z-score FC extract difference (abstinence – satiety) averaged over the R orbitofrontal cortex vs. the difference in the number of EOm for all DEMAND levels of the PFT. The HTPs show a positive relationship between the ΔFC and ΔEOm while the LTPs show a negative relationship for the L Pre : R OFC. **(C)** The z-score FC extract difference (abstinence – satiety) averaged over the L SFG vs. the difference in the number of EOm for all DEMAND levels of the PFT. Both SUBGROUPs show a positive relationship between the ΔFC and ΔEOm for the R v Ins : L SFG. L Pre: Left Precentral gyrus, RvI: Right ventral Insula, R OFC: Right Orbitofrontal Cortex, L SFG: Left Superior Frontal Gyrus, FC: Functional connectivity, EOm: Errors of Omission

## 4. Discussion

The current study employed a ‘dual-stressor’ framework to identify endophenotypic differences in a cohort of active smokers. The cognitive stressor in the form of a Parametric Flanker Task (PFT) identified two smoker SUBGROUPs (i.e., High and Low Task Performers, HTP/LTP) based on their response accuracy on the high DEMAND task condition, independent of nicotine STATEs. However, the imposition of nicotine abstinence as an additional stressor revealed a SUBGROUP*STATE effect of greater sustained attentional lapses (increase in errors of omission, EOm) in the HTPs during abstinence. This behavioral SUBGROUP*STATE interaction was accompanied by neurobiological SUBGROUP*STATE alterations evident in both PFT-evoked activation and network-level interactions within and between ROIs associated with the cognitive construct of attentional control. Taken together, these objectively observed differences at multiple levels of inquiry strongly suggest the presence of valid SUBGROUPs within the smoker population. Such SUBGROUPs may help explain, at least in part, previous inconsistencies in reported cognitive disruptions following acute abstinence and may further inform both foci for and approaches to future targeted treatment interventions.

The current objective characterization of SUBGROUP*STATE effects in early nicotine abstinence have important implications for reconciling uncertainty in the extant literature of abstinence induced cognitive deficits. It is likely that underlying population heterogeneity and compensatory homeostatic mechanisms within smoker cohorts could mitigate the detection of robust abstinence-induced cognitive deficits assumed to arise from neuroplasticity-induced changes with chronic nicotine use (44,45). Specifically, only small to medium effect size deficits have been reported for attention and WM (46), response inhibition and attention and response inhibition (48). Lesage et al.(48,49) also reported no effects on inhibition-related activity, although as seen in the current study, increased EOm in abstinent smokers was observed. In this study, the behavioral performance of the SUBGROUPs identified on the PFT was differentiated by two cognitive components: 1) selective attention (maximizing response accuracy), independent of nicotine STATE and 2) sustained attention (minimizing EOm), which displayed a SUBGROUP*STATE interaction. Electrophysiological indices of cognitive control processes have validated the Flanker paradigm as a modulator of visual attentional control (50,51). Maintaining high accuracy on responded PFT trials requires an effortful allocation of resource-limited, fatigue-prone attentional control to suppress target-irrelevant information (52). In a similar vein, sustained attention is also effortful (53) and the PFT has previously been used to elicit EOm via the temporary depletion of neural control resources (54). Our parametric instantiation of the PFT (25 min. task with a button press required on every trial) thus taxed both attentional control and sustained attention resources.

Based on the above behavioral differences between the SUBGROUPs, the underlying neurobiology described herein help elucidate the mechanisms through which these SUBGROUPs differ. During task performance in the high DEMAND condition, increased activation of multiple brain regions, including the bilateral insula, dorsal anterior cingulate cortex (dACC), right thalamus and the frontoparietal attentional network was observed in the HTPs compared to the LTPs (Fig. 3A, Table S1) across STATE. These brain regions are canonically associated with performance monitoring and sustained attention as observed in contrasts of correct vs incorrect trials in the PFT (55). Further, the dACC and insula are associated with significantly greater BOLD responses for aware vs. unaware errors in a response inhibition task (56). Better performance monitoring through the recruitment of the dACC and insula in the HTPs may in part relate to their better response accuracy. Importantly, these task-evoked effects in the HTPs were independent of nicotine STATE—and thus not related to nicotine withdrawal but potentially indicative of enhanced performance monitoring in the HTP SUBGROUP, while LTPs, who performed at chance levels, showed lower recruitment of these areas.

To elucidate circuits differentially related to attentional control in the two SUBGROUPs we next examined FC pattern differences using the above regions of differential activation as seeds. Across all seeds tested, two identified dyads: L Precentral : R ventral Insula (LPre : RvI) and L posterior Insula : R middle Occipital (LpI : RmO) showed a SUBGROUP*STATE interaction such that only the HTPs demonstrated increased FC during nicotine abstinence between the nodes of the dyads. The RvI and LpI are both components of the SN, which plays a key role in monitoring interoception and regulating body homeostasis (33,57,58). Given the key roles played by LPre, RvI, LpI and RmO in visuospatial attentional control (59,60), the increased FC strength between the *dyad* poles likely contributed to the HTPs ability to selectively attend to high DEMAND PFT trials during abstinence. However, since these regions are also crucial for sustained attention (61,62), the expenditure of attentional resources on selective attention may leave the HTPs susceptible to lapses of sustained attention during nicotine abstinence. The LTPs on the other hand, did not show increases in FC strength between the *dyad* poles and remained relatively impervious to perturbations of sustained attention during nicotine abstinence. Maintenance of attentional control during nicotine abstinence thus may come at the expense of dysregulated sustained attention in the HTPs.

A regression analysis on the four *dyad* poles identified a second set of circuits (Fig 5A) showing a direct relationship between abstinence-induced brain FC changes (ΔFC) and PFT sustained attentional changes (ΔEOm) for the SUBGROUPs. While both SUBGROUPs show a positive relationship between ΔFC and ΔEOm for the RvI : LSFG connection (Fig. 5C), the correlation of ΔFC with ΔEOm appears stronger in the HTPs (vs. LTPs). In a previous study, the LSFG showed increased BOLD response during lapses of attention compared to correct trials in healthy participants performing a Continuous Performance Task (63). The direct positive relationship we observed between the ΔFC and ΔEOm across SUBGROUPs in the LSFG suggests its involvement in suboptimal sustained attention in both SUBGROUPs, but to a greater degree in the HTPs. For the LPre : ROFC connection, the HTPs showed a positive relationship between ΔFC and ΔEOm (Fig. 5B). Among its attributed functions, the ROFC is associated with spatial selective attention (64) and with target detection (65) during visuospatial attention tasks. In light of our findings, the greater involvement of the ROFC during abstinence suggests better attentional control in the HTPs but at the cost of sustained attention. Stronger correlations for the LSFG and ROFC for the HTPs are likely a result of the greater sensitivity to nicotine abstinence in the HTPs, manifest as a greater increase in EOm and increased FC strength in the *dyads* in Fig. 4A.

Although the two stressors (task DEMAND and nicotine STATE) did not interact directly, the attentional demands of the PFT concurrent with the presence/absence of nicotine produced a dynamic break with homeostasis in response to allostatic load (17), revealing differential behavioral (lapses of attention) and neurobiological (network FC) SUBGROUP*STATE interactions, i.e., two distinct SUBGROUPs with variable cognitive abilities. This putative heterogeneity in the smoker endophenotype has been previously implicated through cognitive task response (66), data-driven approaches such as hierarchical clustering on clinical assessment characteristics (67), genetically biased neurobiology (68,69) and cessation treatment outcomes (12).

While it has been hypothesized that the lack of abstinence-induced cognitive effects may be a consequence of small sample sizes and ceiling effects (70), the current sample (n=45), within-subject design and multilevel behavioral and neurobiological analytical strategies reliably demonstrate the underlying heterogeneity within smoker cohorts and compensatory homeostatic mechanisms. This heterogeneity could be a critical independent variable attenuating the detection of abstinence-induced effects on cognitive performance in the extant literature.

While the within-subjects ‘dual stressor’ design revealed objective differences between smoker SUBGROUPs, there are limitations to consider. The PFT as administered is primarily an attentional control task. While it was also able to tax sustained attention, a more direct measurement of sustained attention may unveil further behavioral and neurobiological differences in attentional processes between the SUBGROUPs. Nevertheless, examining both aspects in one task allowed us to look directly at both constructs within the same task. Additionally, as participants were part of a larger treatment study, counterbalancing of the nicotine STATE manipulation was not possible, i.e., the sated scan always preceded the abstinent one and the cohort of smokers were not recruited with the explicit motive of heterogeneous subgrouping.

In sum, by challenging a smoker cohort using concurrent stressors of cognitive DEMAND and nicotine abstinence two SUBGROUPs of smokers were objectively characterized on observed differential susceptibility to abstinence-induced lapses of sustained attention and related underlying differences in task-evoked brain activation and functional network connectivity. The reflection of these SUBGROUP differences at multiple levels of inquiry suggest that this index of smoker heterogeneity may have important clinical utility in predicting smoker susceptibility to NWS-induced cognitive/attentional disruptions, which could lead to triaging NRT to specific subgroups with stronger attentional control but greater abstinence-evoked deficits in sustained attention. Further, the differences in network connectivity may suggest potential differential avenues of treatment intervention via e.g., non-invasive brain stimulation or pharmacological methods and potentially serve as quantitative biomarkers of successful completion of a course of treatment.

## Acknowledgements

This study was supported by the Intramural Research Program of the National Institute on Drug Abuse/National Institutes of Health (NIH) and Food and Drug Administration Grant No. NDA13001-001- 00000 (to EAS). The authors thank Kim Slater, Bridget Moynihan, and Kevin Noemer for assistance with data collection.

## References

1. Allen, S. S., Bade, T., Hatsukami, D. & Center, B. Craving, withdrawal, and smoking urges on days immediately prior to smoking relapse. Nicotine Tob. Res. 10, 35–45 (2008).

2. Snyder, F. R., Davis, F. C. & Henningfield, J. E. The tobacco withdrawal syndrome: performance decrements assessed on a computerized test battery. Drug Alcohol Depend. 23, 259–266 (1989).

3. Kenford, S. L. et al.. Predicting relapse back to smoking: contrasting affective and physical models of dependence. J. Consult. Clin. Psychol. 70, 216–227 (2002).

4. McLaughlin, I., Dani, J. A. & De Biasi, M. Nicotine withdrawal. Curr. Top. Behav. Neurosci. 24, 99–123 (2015).

5. Koob, G. F. & Le Moal, M. Review. Neurobiological mechanisms for opponent motivational processes in addiction. Philos. Trans. R. Soc. Lond. B Biol. Sci. 363, 3113–3123 (2008).

6. Robinson, J. D. et al.. Evaluating the temporal relationships between withdrawal symptoms and smoking relapse. Psychol. Addict. Behav. 33, 105–116 (2019).

7. Piper, M. E. Withdrawal: Expanding a Key Addiction Construct. Nicotine Tob. Res. 17, 1405–1415 (2015).

8. Sheets, E. S., Bujarski, S., Leventhal, A. M. & Ray, L. A. Emotion differentiation and intensity during acute tobacco abstinence: A comparison of heavy and light smokers. Addict. Behav. 47, 70–73 (2015).

9. Perkins, K. A., Briski, J., Fonte, C., Scott, J. & Lerman, C. Severity of tobacco abstinence symptoms varies by time of day. Nicotine Tob. Res. 11, 84–91 (2009).

10. Piper, M. E. et al.. Tobacco withdrawal components and their relations with cessation success. Psychopharmacology 216, 569–578 (2011).

11. Patterson, F. et al.. Toward personalized therapy for smoking cessation: a randomized placebo-controlled trial of bupropion. Clin. Pharmacol. Ther. 84, 320–325 (2008).

12. Lerman, C. et al.. Use of the nicotine metabolite ratio as a genetically informed biomarker of response to nicotine patch or varenicline for smoking cessation: a randomised, double-blind placebo-controlled trial. Lancet Respir Med 3, 131–138 (2015).

13. Benowitz, N., Swan, G., Jacobiii, P., Lessovschlaggar, C. & Tyndale, R. CYP2A6 genotype and the metabolism and disposition kinetics of nicotine. Clinical Pharmacology & Therapeutics vol. 80 457–467 (2006).

14. Fibiger, H. C. Psychiatry, the pharmaceutical industry, and the road to better therapeutics. Schizophr. Bull. 38, 649–650 (2012).

15. Fisher, S. & Reason, J. Handbook of Life Stress, Cognition and Health. (Wiley, 1988).

16. McEwen, B. S. Allostasis and allostatic load: implications for neuropsychopharmacology. Neuropsychopharmacology 22, 108–124 (2000).

17. McEwen, B. S. & Gianaros, P. J. Stress-and allostasis-induced brain plasticity. Annu. Rev. Med. 62, 431– 445 (2011).

18. Richards, J. M. et al.. Biological mechanisms underlying the relationship between stress and smoking: state of the science and directions for future work. Biol. Psychol. 88, 1–12 (2011).

19. Moses, Z. B., Luecken, L. J. & Eason, J. C. Measuring task-related changes in heart rate variability. Conf. Proc. IEEE Eng. Med. Biol. Soc. 2007, 644–647 (2007).

20. McDuff, D., Gontarek, S. & Picard, R. Remote measurement of cognitive stress via heart rate variability. Conf. Proc. IEEE Eng. Med. Biol. Soc. 2014, 2957–2960 (2014).

21. Hendricks, P. S., Ditre, J. W., Drobes, D. J. & Brandon, T. H. The early time course of smoking withdrawal effects. Psychopharmacology 187, 385–396 (2006).

22. Hughes, J. R. Effects of abstinence from tobacco: valid symptoms and time course. Nicotine Tob. Res. 9, 315–327 (2007).

23. Weigard, A., Huang-Pollock, C., Heathcote, A., Hawk, L., Jr & Schlienz, N. J. A cognitive model-based approach to testing mechanistic explanations for neuropsychological decrements during tobacco abstinence. Psychopharmacology 235, 3115–3124 (2018).

24. Harrison, E. L. R., Coppola, S. & McKee, S. A. Nicotine deprivation and trait impulsivity affect smokers’ performance on cognitive tasks of inhibition and attention. Exp. Clin. Psychopharmacol. 17, 91–98 (2009).

25. Nichols, T. T., Gates, K. M., Molenaar, P. C. M. & Wilson, S. J. Greater BOLD activity but more efficient connectivity is associated with better cognitive performance within a sample of nicotine-deprived smokers. Addict. Biol. 19, 931–940 (2014).

26. Jacobsen, L. K. et al.. Effects of smoking and smoking abstinence on cognition in adolescent tobacco smokers. Biol. Psychiatry 57, 56–66 (2005).

27. Mendrek, A. et al.. Working memory in cigarette smokers: comparison to non-smokers and effects of abstinence. Addict. Behav. 31, 833–844 (2006).

28. Ashare, R. L. & Hawk, L. W., Jr. Effects of smoking abstinence on impulsive behavior among smokers high and low in ADHD-like symptoms. Psychopharmacology 219, 537–547 (2012).

29. Patterson, F. et al.. Working memory deficits predict short-term smoking resumption following brief abstinence. Drug Alcohol Depend. 106, 61–64 (2010).

30. Loughead, J. et al.. Working memory-related neural activity predicts future smoking relapse. Neuropsychopharmacology 40, 1311–1320 (2015).

31. Cleck, J. N. & Blendy, J. A. Making a bad thing worse: adverse effects of stress on drug addiction. J. Clin. Invest. 118, 454–461 (2008).

32. Allenby, C. et al.. Brain Marker Links Stress and Nicotine Abstinence. Nicotine Tob. Res. 22, 885–891 (2020).

33. Sutherland, M. T. et al.. Down-regulation of amygdala and insula functional circuits by varenicline and nicotine in abstinent cigarette smokers. Biol. Psychiatry 74, 538–546 (2013).

34. Fedota, J. R. et al.. Nicotine Abstinence Influences the Calculation of Salience in Discrete Insular Circuits. Biol Psychiatry Cogn Neurosci Neuroimaging 3, 150–159 (2018).

35. Fedota, J. R. et al.. Time-Varying Functional Connectivity Decreases as a Function of Acute Nicotine Abstinence. Biol Psychiatry Cogn Neurosci Neuroimaging 6, 459–469 (2021).

36. Fedota, J. R. et al.. Insula Demonstrates a Non-Linear Response to Varying Demand for Cognitive Control and Weaker Resting Connectivity With the Executive Control Network in Smokers. Neuropsychopharmacology 41, 2557–2565 (2016).

37. Forster, S. E., Carter, C. S., Cohen, J. D. & Cho, R. Y. Parametric manipulation of the conflict signal and control-state adaptation. J. Cogn. Neurosci. 23, 923–935 (2011).

38. Welsch, S. K. et al.. Development and validation of the Wisconsin Smoking Withdrawal Scale. Exp. Clin. Psychopharmacol. 7, 354–361 (1999).

39. Heishman, S., Singleton, E. & Pickworth, W. Reliability and validity of a Short Form of the Tobacco Craving Questionnaire. Nicotine & Tobacco Research vol. 10 643–651 (2008).

40. Watson, D., Clark, L. A. & Tellegen, A. Development and validation of brief measures of positive and negative affect: the PANAS scales. J. Pers. Soc. Psychol. 54, 1063–1070 (1988).

41. Cox, R. W. AFNI: software for analysis and visualization of functional magnetic resonance neuroimages. Comput. Biomed. Res. 29, 162–173 (1996).

42. Cox, R. W., Chen, G., Glen, D. R., Reynolds, R. C. & Taylor, P. A. FMRI Clustering in AFNI: False-Positive Rates Redux. Brain Connect. 7, 152–171 (2017).

43. Cole, M. W. et al.. Task activations produce spurious but systematic inflation of task functional connectivity estimates. Neuroimage 189, 1–18 (2019).

44. Ashare, R. L., Falcone, M. & Lerman, C. Cognitive function during nicotine withdrawal: Implications for nicotine dependence treatment. Neuropharmacology 76 Pt B, 581–591 (2014).

45. McClernon, F. J., Addicott, M. A. & Sweitzer, M. M. Smoking abstinence and neurocognition: implications for cessation and relapse. Curr. Top. Behav. Neurosci. 23, 193–227 (2015).

46. Patterson, F. et al.. Varenicline improves mood and cognition during smoking abstinence. Biol. Psychiatry 65, 144–149 (2009).

47. Kozink, R. V., Lutz, A. M., Rose, J. E., Froeliger, B. & McClernon, F. J. Smoking withdrawal shifts the spatiotemporal dynamics of neurocognition. Addict. Biol. 15, 480–490 (2010).

48. Dawkins, L., Powell, J. H., West, R., Powell, J. & Pickering, A. A double-blind placebo-controlled experimental study of nicotine: II--Effects on response inhibition and executive functioning. Psychopharmacology 190, 457–467 (2007).

49. Lesage, E., Sutherland, M. T., Ross, T. J., Salmeron, B. J. & Stein, E. A. Nicotine dependence (trait) and acute nicotinic stimulation (state) modulate attention but not inhibitory control: converging fMRI evidence from Go–Nogo and Flanker tasks. Neuropsychopharmacology vol. 45 857–865 (2020).

50. Maier, M. E., Yeung, N. & Steinhauser, M. Error-related brain activity and adjustments of selective attention following errors. Neuroimage 56, 2339–2347 (2011).

51. McDermott, T. J., Wiesman, A. I., Proskovec, A. L., Heinrichs-Graham, E. & Wilson, T. W. Spatiotemporal oscillatory dynamics of visual selective attention during a flanker task. Neuroimage 156, 277–285 (2017).

52. Faber, L. G., Maurits, N. M. & Lorist, M. M. Mental fatigue affects visual selective attention. PLoS One 7, e48073 (2012).

53. Warm, J. S., Parasuraman, R. & Matthews, G. Vigilance requires hard mental work and is stressful. Hum. Factors 50, 433–441 (2008).

54. Pontifex, M. B., Scudder, M. R., Drollette, E. S. & Hillman, C. H. Fit and vigilant: the relationship between poorer aerobic fitness and failures in sustained attention during preadolescence. Neuropsychology 26, 407–413 (2012).

55. Iannaccone, R. et al.. Conflict monitoring and error processing: New insights from simultaneous EEG– fMRI. NeuroImage vol. 105 395–407 (2015).

56. Orr, C. & Hester, R. Error-related anterior cingulate cortex activity and the prediction of conscious error awareness. Front. Hum. Neurosci. 6, 177 (2012).

57. Craig, A. D. (bud) & (Bud) Craig, A. D. How do you feel — now? The anterior insula and human awareness. Nature Reviews Neuroscience vol. 10 59–70 (2009).

58. Naqvi, N. H. & Bechara, A. The insula and drug addiction: an interoceptive view of pleasure, urges, and decision-making. Brain Struct. Funct. 214, 435–450 (2010).

59. Ikkai, A. & Curtis, C. E. Cortical activity time locked to the shift and maintenance of spatial attention. Cereb. Cortex 18, 1384–1394 (2008).

60. Kim, C., Johnson, N. F. & Gold, B. T. Common and distinct neural mechanisms of attentional switching and response conflict. Brain Res. 1469, 92–102 (2012).

61. Han, S. W. & Marois, R. Functional fractionation of the stimulus-driven attention network. J. Neurosci. 34, 6958–6969 (2014).

62. Nelson, S. M. et al.. Role of the anterior insula in task-level control and focal attention. Brain Struct. Funct. 214, 669–680 (2010).

63. Phillips, R. C., Salo, T. & Carter, C. S. Distinct neural correlates for attention lapses in patients with schizophrenia and healthy participants. Front. Hum. Neurosci. 9, 502 (2015).

64. Shulman, G. L. et al.. Right hemisphere dominance during spatial selective attention and target detection occurs outside the dorsal frontoparietal network. J. Neurosci. 30, 3640–3651 (2010).

65. Luks, T. L., Simpson, G. V., Dale, C. L. & Hough, M. G. Preparatory allocation of attention and adjustments in conflict processing. Neuroimage 35, 949–958 (2007).

66. Sutherland, M. T. et al.. Individual differences in amygdala reactivity following nicotinic receptor stimulation in abstinent smokers. Neuroimage 66, 585–593 (2013).

67. Ding, X. et al.. Evidence of subgroups in smokers as revealed in clinical measures and evaluated by neuroimaging data: a preliminary study. Addiction Biology vol. 24 777–786 (2019).

68. Hong, L. E. et al.. A genetically modulated, intrinsic cingulate circuit supports human nicotine addiction. Proc. Natl. Acad. Sci. U. S. A. 107, 13509–13514 (2010).

69. Li, S., Yang, Y., Hoffmann, E., Tyndale, R. F. & Stein, E. A. CYP2A6 Genetic Variation Alters Striatal-Cingulate Circuits, Network Hubs, and Executive Processing in Smokers. Biol. Psychiatry 81, 554–563 (2017).

70. Rhodes, J. D. & Hawk, L. W., Jr. Smoke and mirrors: The overnight abstinence paradigm as an index of disrupted cognitive function. Psychopharmacology 233, 1395–1404 (2016).

