## Supplementary material for "Not all smokers are alike: The hidden cost of sustained attention during nicotine abstinence"

### Supplementary methods

#### *Participants*

Fifty-nine right-handed participants between 18 and 60 years of age with nicotine dependence were recruited into the study. Exclusion criteria included active drug or alcohol abuse/dependence (other than nicotine), current psychiatric or neurological disorders, and contraindications for magnetic resonance imaging (MRI). Eleven were excluded during final analyses (two for incomplete behavioral data, two for scanning errors, two for abnormal MRI and five for excessive head motion) Overall demographics for the included forty-five participants are described in Table 1. Written informed consent was obtained in accordance with the National Institute on Drug Abuse Intramural Research Program Institutional Review Board.

Upon arrival for their MR scanning sessions, participants completed a medical assessment including a urine test (ASC-11, American Screening Corp., Shreveport, LA) for recent usage of illicit drugs (opiates, oxycodone, benzodiazepines, buprenorphine, cocaine, amphetamines/methamphetamines, tetrahydrocannabinol (THC), methadone, phencyclidine, and methylenedioxymethamphetamine) and an alcohol assessment (Alcohol sensory IV, Intoximeters Inc., St. Louis, MO). Positive urine tests were exclusionary for all drugs except THC. THC-positive urine tests were followed by the Drug Evaluation and Classification neuromotor exam to determine whether the participant was actively intoxicated. Positive neuromotor exams were exclusionary. Nicotine abstinence prior to the second scan was assessed by participant self-report of last cigarette smoked and the measurement of expired carbon monoxide (CO, BreathCO, Vitalograph, Lenexa, KS). An expired CO level of <= 5 ppm was required to physiologically verify nicotine abstinence [1]. In the event of self-reported lapse of abstinence or CO values greater than required level, the abstinence scan was rescheduled.

#### *Behavioral task measures*

Participants underwent functional magnetic resonance imaging (fMRI) while completing a Parametric Flanker Task (PFT) – a modified version of the classic Ericksen flanker task designed to instantiate varying levels of DEMAND for attentional control on a trial-by-trial basis via response conflict. As previously described [2,3], the PFT stimulus array consisted of seven white letters (`S` and/or `H`) displayed horizontally across the center of a black screen. Participants were instructed to respond as quickly and accurately as possible with either their right or left index finger (counterbalanced across participants) corresponding to the target letter in the middle of the string. During congruent trials (no DEMAND), all flankers matched the target letter (SSSSSSS, HHHHHHH). During high DEMAND trials, the target letter differed from all flankers (HHHSHHH, SSSHSSS). In medium DEMAND trials, the target and adjacent flankers were identical with different outermost flankers (HHSSSHH, SSHHHSS). During low DEMAND trials, the target was identical to all but the two outermost flankers (HSSSSSH, SHHHHHS). Runs were composed of a 50/50 split between congruent and incongruent trials (240 trials total: 120 congruent, and 40 per DEMAND condition). Each trial began with the flanker stimuli displayed for 100ms, followed by the target and flankers for 750ms and a jittered inter-trial interval (ITI; 500-7000ms range).

*Subjective measures*

Subjective ratings of withdrawal, craving, and affect were assessed prior to each scanning session. Withdrawal-related symptoms were evaluated via the Wisconsin Smoking Withdrawal Scale (WSWS), a 28-item self-report test comprised of seven subscales (anger, anxiety, concentration, craving, hunger, sadness, sleepiness), and a total score [4]. Craving-related symptoms were evaluated via the short form of the tobacco craving questionnaire (TCQ-SF), a 12-item test comprising four subscales: emotionality, expectancy, compulsivity, and purposefulness [5]. Lastly, affective state was determined via the Positive and Negative Affect Schedule (PANAS), a 20-item test comprising two mood scales which quantify positive and negative affect [6].

*MRI acquisition parameters*

Whole-brain echo planar images were acquired on a 3T Siemens Trio (n=35) and a Prisma scanner (n=10) using a 12-channel head coil. The PFT data, collected in five runs as 39 oblique axial slices (4-mm thick; 30° to anterior commissure–posterior commissure line), were acquired using a T2*-weighted, single-shot gradient echo, echo planar imaging sequence sensitive to blood oxygenation level-dependent (BOLD) effects (720 volumes; repetition time (TR)=2000 ms; echo time (TE)=27 ms; flip angle (FA)=78°; field of view 220 × 220 mm2; image matrix 64 × 64). High-resolution oblique–axial T1-weighted structural images were acquired using a 3D magnetization-prepared rapid gradient-echo (MPRAGE) sequence (TR=1900 ms; TE=3.51 ms; TI=900 ms; FA=9°; voxel size=1 mm3).

*Imaging analysis preprocessing*

Task-evoked pipeline: Initial preprocessing was performed using fMRIprep version 1.5.4 (https://fmriprep.org/en/1.5.4/citing.html). Subsequent preprocessing was performed using AFNI (https://afni.nimh.nih.gov/ version 20191113). Each of the five runs were separately slice-time and motion corrected, detrended and spatially normalized to MNI space via non-linear registration. After motion correction, any two consecutive time points with Euclidean distance derivative values > 0.35 mm were censored [7]. Only participants retaining > 75% volumes in each imaging session after motion censoring were used for further analyses. Framewise displacement (FD) was calculated for each session and the STATE difference (abstinent – sated, ΔFD) was used as a covariate in subsequent analyses. Nuisance covariates, including six-motion parameters were regressed out. Time series were normalized to the percentage of signal change followed by Gaussian spatial smoothing (full width at half-maximum=6 mm).

Task-regressed pipeline: The initial preprocessing for the task-regressed data was identical to that of the task-evoked data described above through the spatial smoothing step. Subsequently, the normalized runs were concatenated, and an intersection of the individual run masks was used to calculate a merge mask. Individual white matter (WM) and cerebrospinal fluid (CSF) masks were obtained by anatomical image segmentation, down-sampled to the EPI resolution, and used to extract signals from non-neuronal sources (*AFNI: 3dfractionize, 3dmaskSVD*).

*FMRIPREP Methods (this section was automatically generated from :https://fmriprep.org/en/1.5.4/citing.html*

Results included in this manuscript come from preprocessing performed using FMRIPREP version 1.5.4[8] RRID:SCR_016216], a Nipype [[9] RRID:SCR_002502] based tool. Each T1w (T1-weighted) volume was corrected for INU (intensity non-uniformity) using N4BiasFieldCorrection v2.1.0 [10] and skull-stripped using antsBrainExtraction.sh v2.1.0 (using the OASIS template). Spatial normalization to the ICBM 152 Nonlinear Asymmetrical template version 2009c [[11], RRID:SCR_008796] was performed through nonlinear registration with the antsRegistration tool of ANTs v2.1.0 [[12], RRID:SCR_004757], using brain-extracted versions of both T1w volume and template. Brain tissue segmentation of cerebrospinal fluid (CSF), white-matter (WM) and gray-matter (GM) was performed on the brain-extracted T1w using fast [[13]] (FSL v5.0.9, RRID:SCR_002823).

Functional data was slice time corrected using 3dTshift from AFNI v16.2.07 [[14], RRID:SCR_005927] and motion corrected using mcflirt (FSL v5.0.9 [[15]]). This was followed by co-registration to the corresponding T1w using boundary-based registration [[16]] with six degrees of freedom, using flirt (FSL). Motion correcting transformations, BOLD-to-T1w transformation and T1w-to-template (MNI) warp were concatenated and applied in a single step using antsApplyTransforms (ANTs v2.1.0) using Lanczos interpolation.

Physiological noise regressors were extracted applying CompCor [[17]]. Principal components were estimated for the two CompCor variants: temporal (tCompCor) and anatomical (aCompCor). A mask to exclude signal with cortical origin was obtained by eroding the brain mask, ensuring it only contained subcortical structures. Six tCompCor components were then calculated including only the top 5% variable voxels within that subcortical mask. For aCompCor, six components were calculated within the intersection of the subcortical mask and the union of CSF and WM masks calculated in T1w space, after their projection to the native space of each functional run. Frame-wise displacement [[18]] was calculated for each functional run using the implementation of Nipype.

Many internal operations of FMRIPREP use Nilearn [[19], RRID:SCR_001362], principally within the BOLD-processing workflow.

For more details of the pipeline see: https://fmriprep.readthedocs.io/en/latest/workflows.html.

**Supplementary results**

*Subjective measurements:* Subjective ratings of withdrawal, craving and affect showed a main effect of STATE. During abstinence, withdrawal, anger (t=2.87, df=40, p=0.006), concentration (t=5.17, df=42, p=6.038e-6) and sleepiness (W=11880, p=2.2e-16, non-parametric Wilcoxon) subscales of the WSWS while sadness showed a trend level effect (t=1.89, df=39, p=0.065). All four subscales of the TCQ-SF craving ratings, emotionality (t=3.61, p=0.00078), expectancy (t=3.74, p=0.00053), compulsivity (t=3.78, p=0.00048), and purposefulness (t=2.32, p=0.025) showed an increase in abstinence. The PANAS showed an increase in negative affect (non-parametric t-test against null; W=2880, p=3.81e-07) and a decrease in positive affect (W=1080, p=3.601e-08) in abstinence. No additional SUBGROUP differences in subjective measures were found for withdrawal (p=0.29), craving (TCQ1 p=0.84, TCQ2 p=0.78, TCQ3 p=0.89, TCQ4 p=0.77), affect (positive affect p=0.67, negative affect p=0.47) or any other SUBGROUP*STATE interaction.

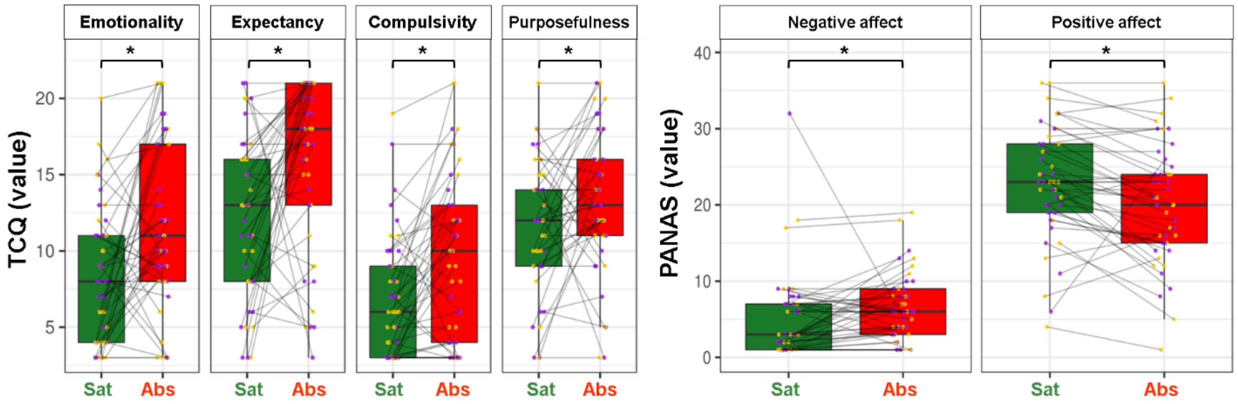

**Figure S1. TCQ and PANAS subjective meeasures:** No SUBGROUP differences or SUBGROUP*STATE interactions for either Tobacco Craving Questionaire (TCQ) or Positive And Negative Affect Schedule (PANAS) were observed. Main effect of STATE was observed with increases in all four subscales of the TCQ, increased negative affect and decreased positive affect during abstinence.

| Voxels | Peak x | Peak y | Peak z | Seed Peak |
|---|---|---|---|---|
| 372 | 53.5 | 33.5 | -2.5 | R Inf Frontal gyrus (p. Orbitalis) |
| 322 | -54.5 | 17.5 | 13.5 | L Inf Frontal gyrus (p. Opercularis) |
| 160 | -52.5 | 19.5 | 25.5 | L Inf Frontal gyrus (p. Triangularis) |
| 112 | 59.5 | -18.5 | 15.5 | R Rolandic operculum |
| 94 | 1.5 | 25.5 | 33.5 | L Mid Cingulate Cortex |
| 79 | 3.5 | 51.5 | 13.5 | R Ant Cingulate Cortex |
| 77 | 51.5 | -14.5 | 45.5 | R Precentral gyrus |
| 66 | 13.5 | -10.5 | 7.5 | R Thalamus |
| 63 | 47.5 | 27.5 | 21.5 | R Inf Frontal gyrus (p. Triangularis) |
| 46 | -4.5 | 43.5 | 27.5 | L Sup Medial gyrus |
| 43 | -62.5 | -56.5 | 23.5 | L Supramarginal gyrus |
| 37 | -56.5 | -4.5 | 23.5 | L Postcentral gyrus |
| **37** | **-44.5** | **1.5** | **35.5** | **L Precentral gyrus** |
| 31 | -56.5 | -52.5 | 31.5 | L Supramarginal gyrus |
| 28 | -54.5 | -32.5 | -6.5 | L Mid Temporal gyrus |
| 28 | 41.5 | -8.5 | -6.5 | R Insula |
| 28 | -14.5 | -78.5 | -0.5 | L Lingual gyrus |
| **27** | **-36.5** | **-10.5** | **11.5** | **L Insula** |
| 24 | 37.5 | -14.5 | 43.5 | R Precentral gyrus |

**Table S1. Parametric Flanker Task evoked BOLD response clusters for the high DEMAND responded trials:** Clusters with differential activation for the HTP compared to the LTP SUBGROUP for the correct – errors of commission trials contrast at the high DEMAND condition across SESSION. These clusters were subsequently used as seeds in a whole-brain seed-based functional connectivity analysis. * denotes the peak for the cluster.

Coordinates are in MNI space, order = LPI; CM, center of mass of the voxel cluster; L, left; R, right; Inf, inferior; Ant, anterior; Mid, middle; SMA, supplementary motor area; Sup, superior; p., pars, voxel size=8mm$^3$. Bolded regions are the seeds that formed a dyad in the whole-brain analysis.

| | | Voxels | Peak x | Peak y | Peak z |
|---|---|---|---|---|---|
| **Dyad1** | **Seed:** L Precentral gyrus | 37 | -44.5 | 1.5 | 35.5 |
| | **Target:** R Vent Insula | 81 | 43.5 | 9.5 | -8.5 |

| | | | | | |
|---|---|---|---|---|---|
| **Dyad2** | **Seed:** L Pos Insula | 27 | -36.5 | -10.5 | 11.5 |
| | **Target:** R Mid Occipital gyrus | 75 | 51.5 | -76.5 | 3.5 |

**Table S2.** Functional connectivity dyad cluster information during PFT-regressed fMRI (Figure 4): *dyad1* with the L Precentral seed and the R vent Insula "target" and *dyad2* with the L pos Insula seed and the R mid Occipital "target". A "target" in our analysis was defined as a significant cluster arising from a whole-brain seed-based FC analysis with SUBGROUP as a between-subjects factor and STATE as a within-subjects factor.

coordinate order = LPI, CM = center of mass of the voxel cluster, L = left, R = right, Vent = ventral, Pos = posterior, Mid = middle, voxel size=8mm$^3$

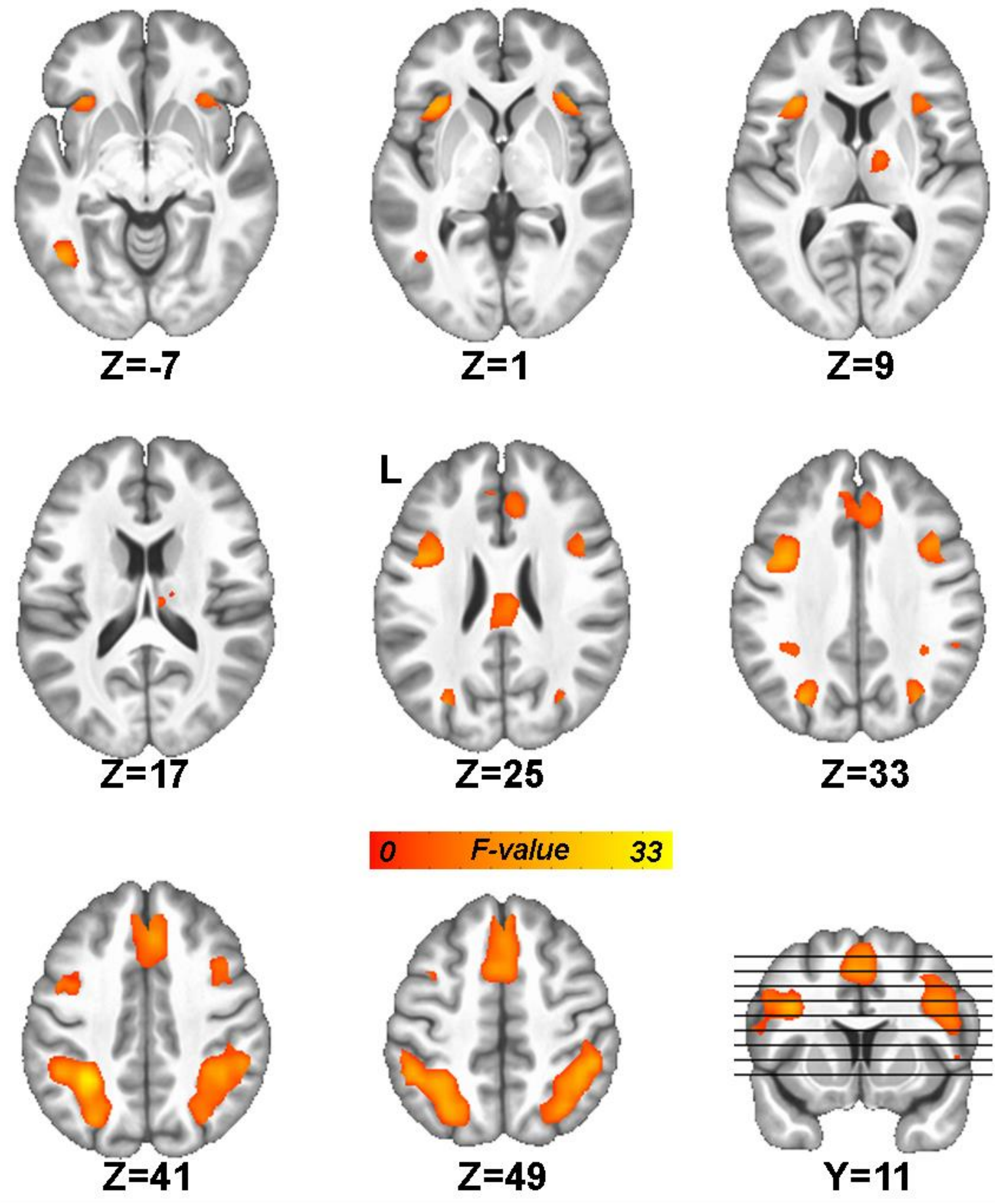

**Figure S2. Parametric Flanker Task Map.** Main Effect of Difficulty (Number of Flankers). PFT strongly activated attentional control and salience network regions, including bilateral insula and anterior cingulate cortex. Whole-brain corrected (p-voxel < 0.001, FWE α < 0.05). Coordinates are in MNI space, order = LPI, L = left

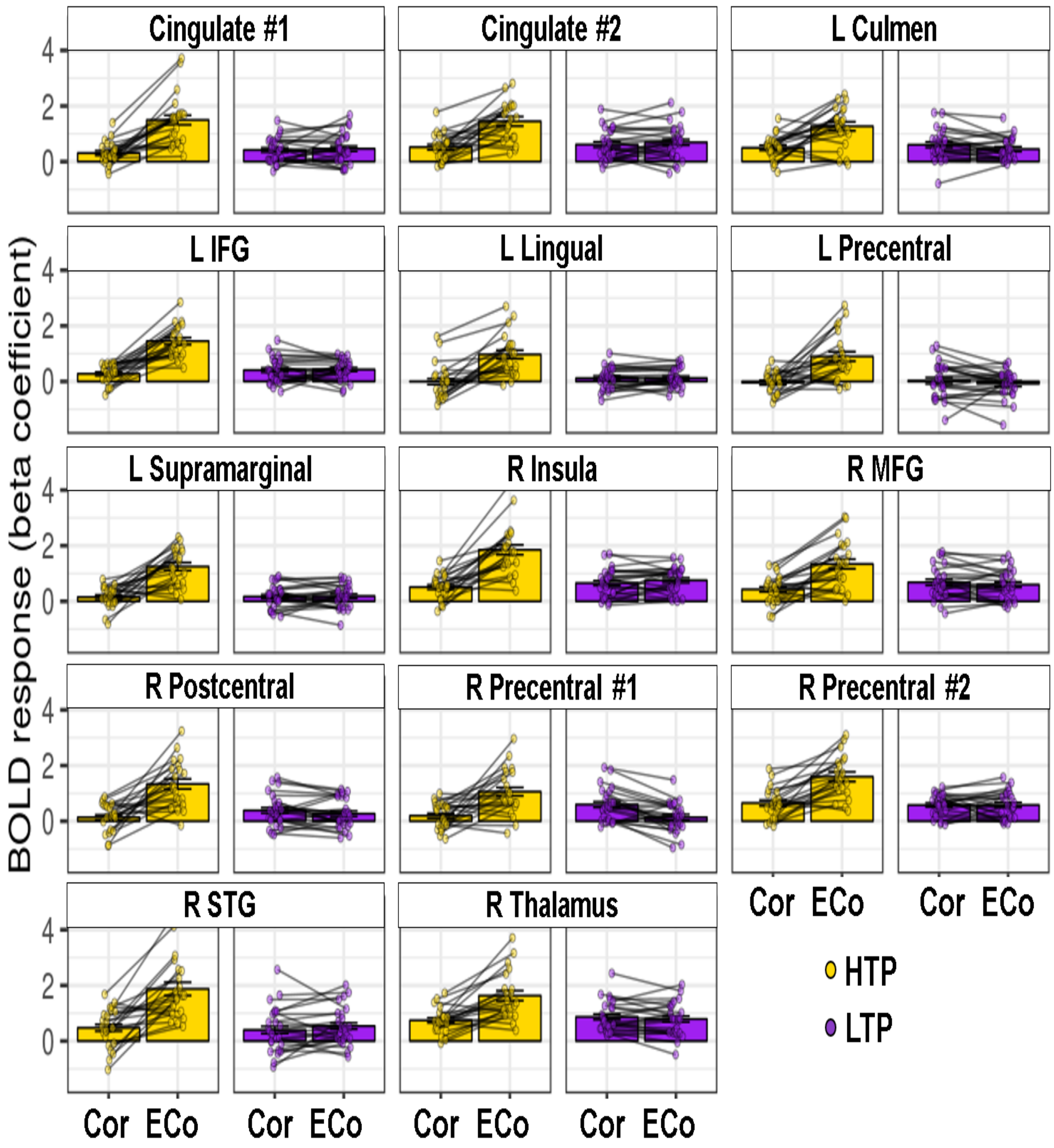

**Figure S3. Parametric Flanker Task extracted BOLD response for the high DEMAND responded trials:** Extracted β coefficients from the clusters in Fig 3A (averaged across SESSION) for corrects (Cor) and errors of commission (ECo) in the HTP and LTP (High Task Performers and Low Task Performers) SUBGROUPs. *IFG: inferior frontal gyrus, MFG: middle frontal gyrus, STG: superior temporal gyrus*

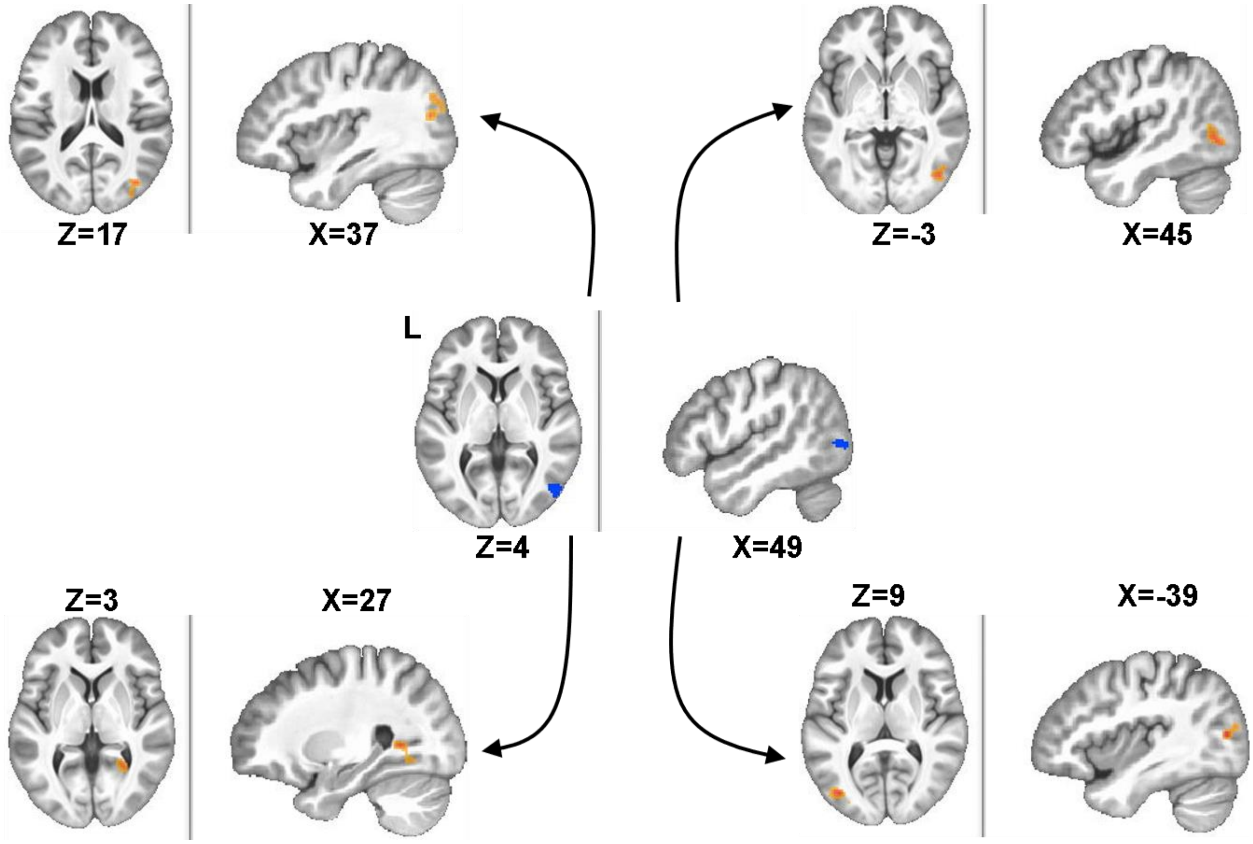

**Figure S4. SUBGROUP differences in the relation of abstinence-induced changes in whole-brain FC (ΔFC) to abstinence-induced changes in sustained attention (ΔEOm).** Whole-brain regression analyses starting with the seed at the R middle occipital (central, blue) leads to “targets” (orange) in the R middle occipital (top left), R inferior occipital (top right). R lingual (bottom left) and L middle occipital (bottom right). The arrows do not imply directionality or causality. Coordinates are in MNI space, order = LPI, L = left

| Seed | Voxels | Peak x | Peak y | Peak z | Regression region | Voxels | Peak x | Peak y | Peak z |
|---|---|---|---|---|---|---|---|---|---|
| L Precentral | 37 | -44.5 | 1.5 | 35.5 | R Inf Frontal gyrus (p. Orbitalis) | 120 | 29.5 | 37.5 | -18.5 |
| R Insula | 81 | 43.5 | 9.5 | -8.5 | L Sup Frontal | 102 | -14.5 | 67.5 | 5.5 |
| R Mid Occipital | 75 | 51.5 | -76.5 | 3.5 | R Mid Occipital | 186 | 37.5 | -76.5 | 17.5 |
| | | | | | R Inferior Occipital | 167 | 45.5 | -72.5 | -2.5 |
| | | | | | R Lingual gyrus | 144 | 27.5 | -54.5 | 3.5 |
| | | | | | L Mid Occipital | 128 | -38.5 | -78.5 | 9.5 |

**Table S3. Clusters of brain (ΔFC) and behavior (ΔEOm) regression between SUBGROUP STATE differences (abstinence – satiety) using individual Z-scored FC extracts (averaged over “target” ROI) and EOm overall DEMAND levels of the PFT**.

coordinate order = LPI; CM, center of mass of the voxel cluster; L, left; R, right; Inf, inferior; Ant, anterior; Mid, middle; SMA, supplementary motor area; Sup, superior; p., pars

**References**

1. Javors, M. A., Hatch, J. P. & Lamb, R. J. Cut-off levels for breath carbon monoxide as a marker for cigarette smoking. *Addiction* **100**, 159–167 (2005).
2. Forster, S. E., Carter, C. S., Cohen, J. D. & Cho, R. Y. Parametric manipulation of the conflict signal and control-state adaptation. *J. Cogn. Neurosci.* **23**, 923–935 (2011).
3. Fedota, J. R. *et al.* Insula Demonstrates a Non-Linear Response to Varying Demand for Cognitive Control and Weaker Resting Connectivity With the Executive Control Network in Smokers. *Neuropsychopharmacology* **41**, 2557–2565 (2016).
4. Welsch, S. K. *et al.* Development and validation of the Wisconsin Smoking Withdrawal Scale. *Exp. Clin. Psychopharmacol.* **7**, 354–361 (1999).
5. Heishman, S., Singleton, E. & Pickworth, W. Reliability and validity of a Short Form of the Tobacco Craving Questionnaire. *Nicotine & Tobacco Research* vol. 10 643–651 (2008).
6. Watson, D., Clark, L. A. & Tellegen, A. Development and validation of brief measures of positive and negative affect: the PANAS scales. *J. Pers. Soc. Psychol.* **54**, 1063–1070 (1988).
7. Power, J. D., Barnes, K. A., Snyder, A. Z., Schlaggar, B. L. & Petersen, S. E. Spurious but systematic correlations in functional connectivity MRI networks arise from subject motion. *Neuroimage* **59**, 2142–2154 (2012).
8. Esteban, O. *et al.* fMRIPrep: a robust preprocessing pipeline for functional MRI. *Nat. Methods* **16**, 111–116 (2019).
9. Gorgolewski, K. *et al.* Nipype: a flexible, lightweight and extensible neuroimaging data processing framework in python. *Front. Neuroinform.* **5**, 13 (2011).
10. Tustison, N. J. *et al.* N4ITK: improved N3 bias correction. *IEEE Trans. Med. Imaging* **29**, 1310–1320 (2010).

11. Fonov, V. S., Evans, A. C., McKinstry, R. C., Almli, C. R. & Collins, D. L. Unbiased nonlinear average age-appropriate brain templates from birth to adulthood. *NeuroImage* vol. 47 S102 (2009).

12. Avants, B. B., Epstein, C. L., Grossman, M. & Gee, J. C. Symmetric diffeomorphic image registration with cross-correlation: evaluating automated labeling of elderly and neurodegenerative brain. *Med. Image Anal.* **12**, 26–41 (2008).

13. Zhang, Y., Brady, M. & Smith, S. Segmentation of brain MR images through a hidden Markov random field model and the expectation-maximization algorithm. *IEEE Trans. Med. Imaging* **20**, 45–57 (2001).

14. Cox, R. W. AFNI: software for analysis and visualization of functional magnetic resonance neuroimages. *Comput. Biomed. Res.* **29**, 162–173 (1996).

15. Jenkinson, M., Bannister, P., Brady, M. & Smith, S. Improved optimization for the robust and accurate linear registration and motion correction of brain images. *Neuroimage* **17**, 825–841 (2002).

16. Greve, D. N. & Fischl, B. Accurate and robust brain image alignment using boundary-based registration. *Neuroimage* **48**, 63–72 (2009).

17. Behzadi, Y., Restom, K., Liau, J. & Liu, T. T. A component based noise correction method (CompCor) for BOLD and perfusion based fMRI. *Neuroimage* **37**, 90–101 (2007).

18. Power, J. D. *et al.* Methods to detect, characterize, and remove motion artifact in resting state fMRI. *Neuroimage* **84**, 320–341 (2014).

19. Abraham, A. *et al.* Machine learning for neuroimaging with scikit-learn. *Front. Neuroinform.* **8**, 14 (2014).
